# Flowers are leakier than leaves but cheaper to build

**DOI:** 10.1101/2023.04.11.536372

**Authors:** Adam B. Roddy, C. Matt Guilliams, Paul V.A. Fine, Stefania Mambelli, Todd E. Dawson, Kevin A. Simonin

## Abstract

- Producing and maintaining flowers is essential for reproduction in most angiosperms, underpinning population persistence and speciation. Although the physiological costs of flowers often oppose pollinator selection, these physiological costs have rarely been quantified.
- We measured a suite of physiological traits quantifying the water and carbon costs and drought tolerance on flowers and leaves of over 100 phylogenetically diverse species, including water and dry mass contents, minimum epidermal conductance to water vapor (*g*_*min*_), vein density, and dry mass per area.
- Although there was substantial variation among species, flowers had significantly higher *g*_*min*_ and water content per unit area than leaves, but significantly lower vein density and dry mass per area than leaves. Both leaves and flowers exhibited similarly strong scaling between dry mass investment and water content.
- The higher *g*_*min*_ of flowers offset their higher water content, suggesting that flowers may desiccate more rapidly than leaves during drought. The coordination between dry mass and water investment suggests that flowers rely on a hydrostatic skeleton to remain upright rather than on a carbon-based skeleton. For short-lived structures like flowers, water may be relatively cheaper than carbon, particularly given the relatively high amount of water loss per unit of carbon synthesized in photosynthesis.

## Main text

Flowers are critical to reproduction in angiosperms and have been credited with promoting diversification and the rapid spread of flowering plants globally (Sanderson & Donoghue, 1994; Crepet & Niklas, 2009; Leslie *et al*., 2021). Although they are typically short-lived, flowers require resources, such as carbon, water, and nutrients, for their production and maintenance (Reekie & Bazzaz, 1987a,b; Ashman & Schoen, 1994; Song *et al*., 2022). Water, in particular, is needed throughout development and anthesis for a variety of functions, including driving growth and expansion, keeping flowers turgid and on display for pollinators, providing rewards such as nectar, and for regulating temperature (Bazzaz *et al*., 1987; Galen *et al*., 1999; Patiño & Grace, 2002; Chapotin *et al*., 2003; De la Barrera & Nobel, 2004; Roddy & Dawson, 2012; Roddy, 2019; Treado *et al*., 2022). Additionally, flowers regularly lose water to the atmosphere, and this water loss may increase during hot and dry conditions often associated with droughts (Hew *et al*., 1980; Feild *et al*., 2009; Teixido & Valladares, 2014; Sinha *et al*., 2022). Flower water balance is, therefore, critical to flower function, yet surprisingly little is known about the mechanisms of water balance in flowers, how physiological traits related to water and carbon influence the costs of floral display, and how floral hydraulic traits affect drought responses (Roddy *et al*., 2016; Bourbia *et al*., 2020; Roddy *et al*., 2021; McMann *et al*., 2022).

The rate of water loss from flowers–and, indeed, from all aerial organs of plants–is ultimately determined by the atmospheric conditions that drive the net loss of water from the plant to atmosphere (e.g. solar radiation, temperature, humidity, windspeed) and by the structure of the epidermis, which controls the total cuticular surface conductance to water vapor (*g*_*t*_). Stomata in the epidermis are the primary pathway for water movement from plants to the atmosphere, and their sizes and densities influence maximum rates of transpirational water loss (Hetherington & Woodward, 2003; Franks & Beerling, 2009). Compared to leaves, flowers often have relatively few, if any, stomata on their petals and petaloid structures (Hew *et al*., 1980; Lipayeva, 1989; Roddy *et al*., 2016; Zhang *et al*., 2018). Under well-watered conditions, the high densities of stomata on angiosperm leaves allow transpiration rates from leaves to exceed those of flowers (Feild *et al*., 2009; Roddy *et al*., 2018). However, under drought conditions, leaf stomata close to limit water loss, causing any remaining water vapor flux to be due to the minimum epidermal surface conductance (*g*_*min*_), which is due to the conductance of the cuticle and any incompletely closed stomata (Kerstiens, 1996; Duursma *et al*., 2019; Márquez *et al*., 2022). After drought induced stomatal closure in leaves, water loss from flowers can be as high as or even exceed water loss from leaves (Sinha *et al*., 2022), suggesting that corolla *g*_*min*_ may hinder the ability of plants to maintain floral display during periods of water stress (Lambrecht, 2013; Buschhaus *et al*., 2015; Bourbia *et al*., 2020). Yet, despite the influence of *g*_*min*_ on flower and whole plant hydration and its role in regulating flower temperature, *g*_*min*_ has been measured on flowers of only a few species (Patiño & Grace, 2002; Roddy *et al*., 2016; Roddy, 2019; Bourbia *et al*., 2020).

Here we compared flowers and leaves in a set of physiological traits that influence water balance, particularly during drought (Brodribb *et al*., 2007; Boyce *et al*., 2009; Simonin *et al*., 2013; Roddy *et al*., 2016; Roddy *et al*., 2018; Duursma *et al*., 2019; Bourbia *et al*., 2020). We measured *g*_*min*_, water content per unit projected surface area (*W*_*area*_) and dry mass (*W*_*mass*_), vein density (*D*_*v*_), and dry mass per area (leaf mass per area, *LMA*, or petal mass per area, *PMA*) of flowers and leaves for over 100 species from 41 angiosperm families growing in a common garden to determine how these physiological traits differ among organs and influence the costs of floral construction and maintenance.

Flower petals differed significantly in most of the water balance traits we evaluated (Figure 1; Table 1). Water content per unit dry mass (*W*_*mass*_), which is positively related to hydraulic capacitance (Ogburn & Edwards, 2012; Roddy *et al*., 2019), was significantly higher in petals than in leaves. Flower petals also had higher *W*_*area*_ than leaves, though this difference was not significant after accounting for shared evolutionary history, and the range of *W*_*area*_ among species was larger for petals than it was for leaves. With a mean of 12.68 mmol m^-2^ s^-1^, flowers had significantly higher *g*_*min*_ than their neighboring leaves. Leaf *g*_*min*_ had a mean of 4.65 mmol m^-2^ s^-1^, which was similar to the interspecific mean of a recent literature survey (mean of 4.9 mmol m^-2^ s^-1^) (Duursma *et al*., 2019) and similar to some tropical leaves (Slot *et al*., 2021). Of the 101 species for which there were *g*_*min*_ data for both leaves and flowers, only 27 species had flowers with lower *g*_*min*_ than leaves.

**Table 1.** Pairwise differences in traits between leaves and flowers.

| trait | mean,<br>leaf | mean, corolla | non-phylogenetic |  | phylogenetic |  |
| --- | --- | --- | --- | --- | --- | --- |
|  |  |  | t (df) | P | t (df) | P |
| $W_{area}$ (mol m<sup>-2</sup> s<sup>-1</sup>) | 8.01 | 9.72 | 2.17 (103) | 0.03 | 1.33 (101) | 0.19 |
| $W_{mass}$ (g g<sup>-1</sup>) | 2.93 | 6.64 | 13.54 (103) | < 0.0001 | 5.16 (101) | < 0.0001 |
| $g_{min}$ (mmol m<sup>-2</sup> s<sup>-1</sup>) | 4.65 | 12.68 | 7.71 (100) | < 0.0001 | 4.50 (98) | < 0.0001 |
| $Dv$ (mm mm<sup>-1</sup>) | 5.66 | 2.57 | 13.23 (85) | < 0.0001 | 5.60 (83) | < 0.0001 |
| $\tau_{VPD=1}$ (hr) | 47.83 | 21.86 | 8.21 (98) | < 0.0001 | 4.73 (96) | < 0.0001 |
| LMA or PMA (g m<sup>-2</sup>) | 51.01 | 27.09 | 7.29 (106) | < 0.0001 | 2.29 (104) | 0.02 |

**Figure 1.**
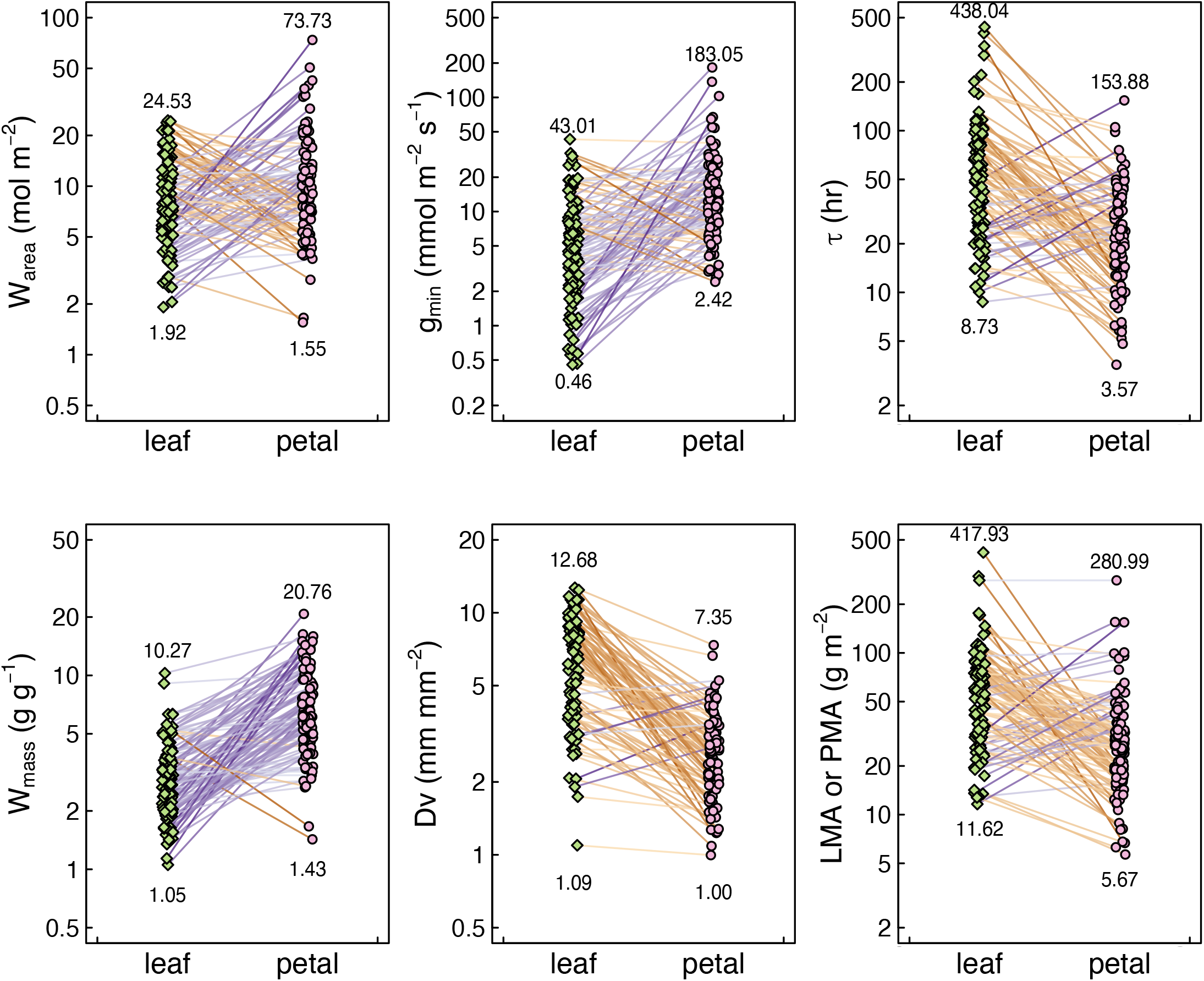
Pairwise difference in trait values between leaves (green diamonds) and petals (pink circles). Lines connect points of the same species, and connecting lines are colored according to the magnitude of the difference in trait values between organs with orange colors indicating leaves have a higher trait value and purple colors indicating flowers have a higher trait value. Numbers above and below point clouds indicate the organ-specific maximum and minimum trait values, i.e. the interspecific range of trait values.

**Figure 2.**
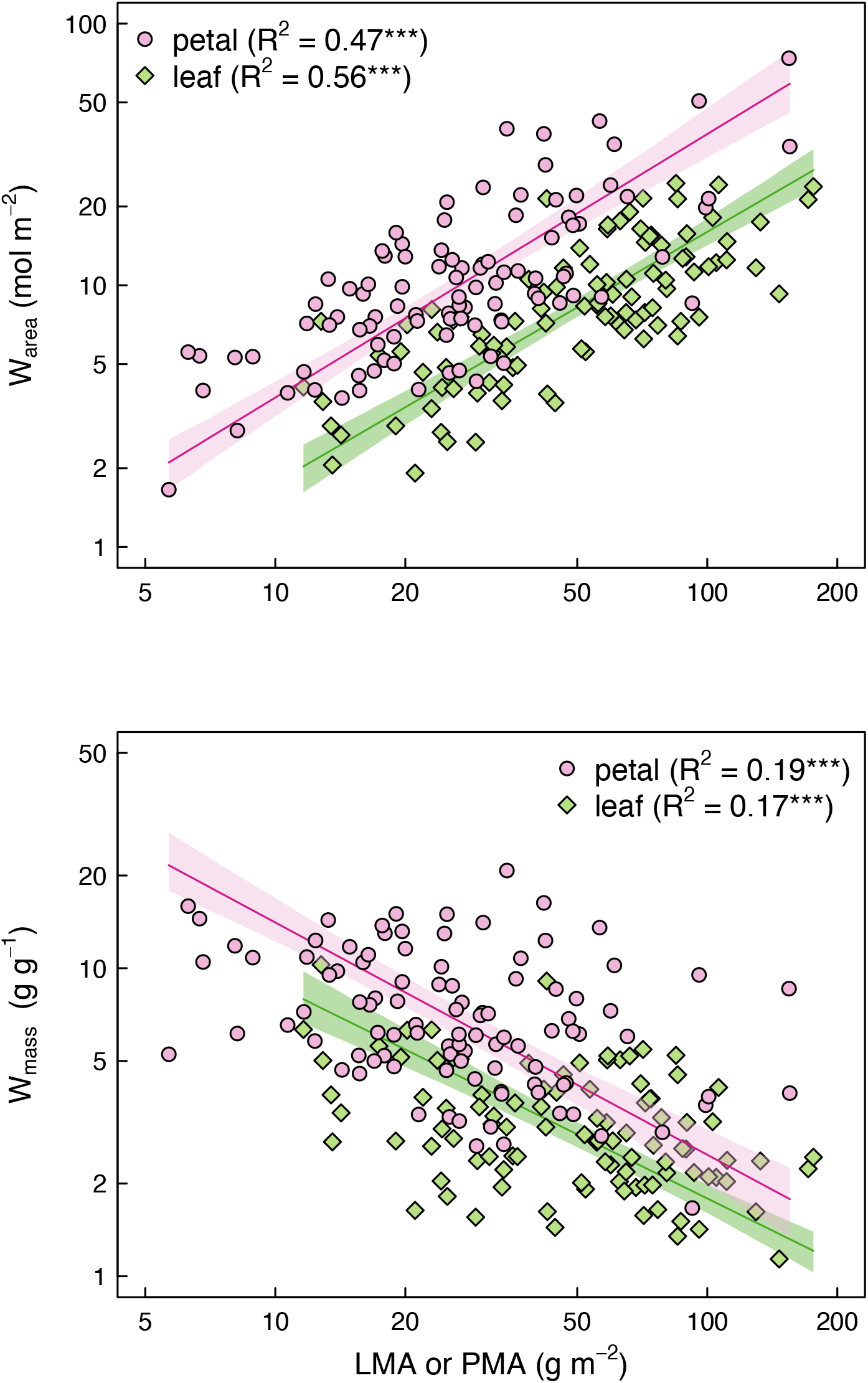
Leaf or petal mass per area (*LMA* or *PMA*, respectively) are coordinated with water content calculated (a) per unit area and (b) per unit dry mass. Pink and green lines and shading represent the standard major axis regressions and 95% confidence intervals.

The relatively high *g*_*min*_ of flowers highlights that flowers may contribute significantly to whole-plant water budgets during flowering periods (Lambrecht & Dawson, 2007; Lambrecht, 2013). In many species, flowers are positioned distal to leaves, resulting in leaf shading and suppressing foliar transpiration. Additionally, because flowers are often positioned in the hottest, driest parts of the plant crown, their high *g*_*min*_ may translate into high rates of water loss (Roddy & Dawson, 2012). For example, previous work on avocado has shown that a combination of high flower transpiration and high total flower surface area resulted in flowers accounting for ∼13% of total canopy water loss (Whiley *et al*., 1988). Given that corolla *g*_*min*_ is, on average, higher than leaf *g*_*min*_ (Figure 1), flower water loss can be as high as or even exceed water loss from leaves and potentially dominate total canopy transpiration (Lambrecht, 2013). More work is needed at the whole-plant scale to characterize whether and when flower water loss can detrimentally impact whole-plant water status and potentially precipitate hydraulic failure in vegetative organs (Nobel, 1977; Galen *et al*., 1999; Lambrecht & Dawson, 2007; Zhang & Brodribb, 2017; Bourbia *et al*., 2020).

Using our measurements of *g*_*min*_ and *W*_*area*_ we calculated water residence times (τ) for leaves and flowers, assuming a constant vapor pressure deficit (VPD) of 1 kPa. Despite having slightly higher *W*_*area*_, flowers had significantly shorter τ than leaves, driven by their higher *g*_*min*_ (Figure 1; Table 1). A shorter τ among flowers suggests that when water loss exceeds water supply, e.g. during drought, flowers would desiccate more rapidly than leaves due to their higher *g*_*min*_. Under higher VPD than the mild 1 kPa we used, τ would be even shorter. For example, increasing VPD to 2.5 kPa reduces the average τ for flowers from 21.86 hr to 8.74 hr and that for leaves from 47.83 hr to 19.13 hr. Under this scenario, the average flower would desiccate within a day without new water supply, consistent with previous reports for *Calycanthus* flowers during a heatwave (Roddy *et al*., 2018).

Despite having higher water contents and hydraulic capacitance (Fig. 1; Roddy *et al*. (2019)), high *g*_*min*_ may require that flowers have constant supplies of water to remain turgid. That *g*_*min*_ has been shown to scale with whole-flower hydraulic conductance reiterates the role of *g*_*min*_ in regulating flower water balance (Roddy *et al*., 2016). Despite early suggestions that flowers may be hydrated primarily by the phloem (Trolinder *et al*., 1993; Chapotin *et al*., 2003), the phloem may not contribute meaningful amounts of water to the overall flower water budget (Feild *et al*., 2009; Roddy *et al*., 2018; McMann *et al*., 2022), and given a relatively shorter intrinsic τ flowers may need to remain connected to the xylem hydraulic system to avoid desiccation (Feild *et al*., 2009; Roddy *et al*., 2018). In leaves and flowers, liquid water is delivered primarily by the network of veins that traverse the leaf and petal. However, flower petals had significantly lower *D*_*v*_ than leaves [Figure 1; Table 1; Roddy *et al*. (2013); Zhang *et al*. (2018)], suggesting that flowers have a lower hydraulic conductance than leaves(Brodribb *et al*., 2007; Roddy *et al*., 2016). The lower *D*_*v*_ despite higher *g*_*min*_ reiterates that flowers may be particularly vulnerable to drought, when water loss may outpace hydraulic supply, leading to rapid declines in flower water potential that could cause declines in stem water potential (Bourbia *et al*., 2020). However, the higher *W*_*area*_ and higher hydraulic capacitance of flowers (i.e. high *W*_*mass*_) would minimize changes in water potential despite their having a low hydraulic conductance. The high hydraulic capacitance of flowers would, therefore, suppress diurnal variation in corolla water potential, reducing the impact of excessive floral water loss on the water potentials of hydraulically upstream organs and reducing the likelihood that flower water potential would decline enough to cause embolism spread (Zhang & Brodribb, 2017; Roddy *et al*., 2018; Roddy *et al*., 2019).

Understanding how *g*_*min*_, *W*_*area*_, and *W*_*mass*_ of flowers interact with hydraulic traits of leaves and stems, and by extension whole plant water and carbon balance, will be critical to better characterizing plant responses to changes in water availability.

Although they had higher water contents than leaves, petals had significantly lower dry mass per area than leaves (Figure 1; Table 1), which has implications for the biomechanics of flower petals. For short-lived structures like flower petals, reducing the costs of floral display has likely been favored by selection (Roddy *et al*., 2016; Olson & Pittermann, 2019; Roddy, 2019). Longer-lived flowers may incur higher carbon costs because they may need to withstand attack by floral enemies (Ashman, 1994; Roddy *et al*., 2021; Boaventura *et al*., 2022; Song *et al*., 2022). How biomass costs of flowers are related to other resources can be variable and context-dependent (Bazzaz *et al*., 1987; Reekie & Bazzaz, 1987b; Roddy *et al*., 2021), but the water and carbon costs may be coupled in important ways. Supplying more water to flowers would require a denser networks of veins, which are carbon-rich and potentially costly to produce. Similarly, better limiting water loss by building thicker or denser cuticles could also require more carbon investment (Buschhaus *et al*., 2015; Cheng *et al*., 2019). Yet, because every molecule of carbon requires at least 400 molecules of water to be transpired (Nobel *et al*., 2005), for short-lived structures such as flowers, water may be relatively cheaper than carbon, suggesting that flowers may employ a hydrostatic skeleton rather than an expensive, carbon-based skeleton for structural support. We tested whether floral display is cheaper in terms of carbon due to higher initial investment of water by examining the relationship between water content and dry mass investment. *W*_*area*_ scaled positively with *LMA* (slope = 0.96 [0.84, 1.09], R^2^ = 0.56, P < 0.0001) and *PMA* (slope = 1.01 [0.88, 1.16], R^2^ = 0.52, P < 0.0001), with statistically indistinguishable slopes between organs (P = 0.47). However, flowers had a significantly higher intercept to this scaling relationship (t = 2.69, df = 98, P < 0.01). Similarly, *W*_*mass*_ scaled negatively with both *PMA* (slope = -0.75 [-0.91, -0.63], R^2^ = 0.14, P < 0.001) and *LMA* (slope = -0.69 [-0.83, -0.58], R^2^ = 0.16, P < 0.0001) with slopes indistinguishable between organs (P = 0.36) but a significantly higher intercept among flowers (t = 2.57, df = 98, P < 0.05). The higher intercepts in these scaling relationships support the hypothesis that flowers rely on a hydrostatic skeleton maintained by high water content and cheap, thin cell walls that allow for relatively low dry mass per unit area. This combination of traits–high water content and low dry mass–would also explain why flowers have higher hydraulic capacitance than leaves, as thin cell walls with a low modulus of elasticity would allow large changes in cell volume with relatively small changes in water potential (Roddy *et al*., 2019).

While there are numerous implications of higher *g*_*min*_ in flowers, it is important to consider why *g*_*min*_ is higher in flowers. We propose two alternative explanations for this pattern. First, if reducing *g*_*min*_ requires greater carbon investment (e.g. through additional cuticular waxes) (Cheng *et al*., 2019), then the additional carbon required to further reduce water loss may not be worth paying in such a short-lived organ. Second, cuticle structure and composition may experience divergent selection for multiple functions. For the majority of angiosperm species, petaloid organs attract pollinators, with one mechanism of attraction being the release of volatile organic compounds (Dudareva *et al*., 2013). Corolla cuticles are structurally different from those of neighboring leaves (Jetter *et al*., 2008; Cheng *et al*., 2019), and cuticle structure can influence both volatile emission and conductance to water vapor (Goodwin *et al*., 2003; Buschhaus *et al*., 2015; Liao *et al*., 2021). Thus, pollinator selection on volatile organic compound emissions may be tightly coupled to the mechanisms of flower water balance. Or, water loss from the flower may itself be used as a signal for pollinators (Dahake *et al*., 2022). Regardless of which explanation may be true, both mechanisms suggest that climate change may alter the selection dynamics on floral hydraulic traits. Both increased likelihood of drought and pollinator declines could shift the relative costs of producing and maintaining flowers (Thomann *et al*., 2013; Gallagher & Campbell, 2017; Kuppler & Kotowska, 2021). Understanding how shifting selective regimes may impact floral function and evolution will be important in understanding the future viability of flowering plants globally.

## Acknowledgments

This work was supported by a grant from the Jepson Herbarium of the University of California, Berkeley. ABR was supported by a US NSF Graduate Research Fellowship and by NSF grant CMMI-2029756. S.F. Oberbauer and D.C. Paiva provided valuable feedback on an earlier draft. We thank H. Forbes at the University of California Botanic Garden for facilitating access to plant material.

## Author contributions

ABR, CMG, PVAF, SM, TED, and KAS conceptualized the study. ABR and CMG collected the data. ABR and KAS analyzed the data. ABR wrote the manuscript, and all authors edited the manuscript.

## Materials and Methods

### Plant material

Plants were sampled in 2011 and 2012 at the University of California Botanical Garden in Berkeley, CA, USA, where plants are maintained well-watered throughout the year. The species samples represent a phylogenetically and ecologically diverse set of species that vary in stature, habitat, and flowering phenology (Table S1). We sampled in the morning, between 8-9 am, by excising a flowering shoot and immediately recutting the cut end under water a few inches apical to the initial cut or, for large woody species, one node more apical from the initial cut. If leaves were not present on this shoot, we excised a leafy shoot in a similar way. Cut ends remained in water, and the shoots were kept in a bucket to shield them from desiccation during transport. Shoots were transported back to the laboratory within 1 hour and kept in the dark prior to trait sampling. We measured leaves and flowers on the same plant and, for most species, we sampled one individual plant per species. While this sampling approach may not be used to characterize average trait values for each species, it is particularly well-suited to test whether organs differ in traits

### Trait measurements

Minimum surface conductance (*g*_*min*_) was measured by excising a leaf at the petiole basis or a flower petal. When flower petals were fused, we excised all fused petals, keeping them intact to minimize cut surfaces. Immediately upon excision, the cut surfaces of the petals or leaf petiole were sealed with either cyanoacrylate glue or petroleum jelly. Samples were either hung or placed on a mesh screen in a dark cabinet with a fan blowing directly on them in order to maximize the boundary layer conductance. Every 5-20 minutes, samples were weighed on an analytical balance (Sartorius CPA225D, resolution = 0.0001 g) and the temperature and humidity inside the chamber recorded. After approximately 10 measurements, each sample was either scanned or photographed for subsequent measurement of its area and then dried at 70^°^C for at least 72 hours for subsequent dry mass measurement. *g*_*min*_ was calculated from measurements of mass, temperature, and humidity using the spreadsheet available https://prometheusprotocols.net/function/gas-exchange-and-chlorophyll-fluorescence/stomatal-and-non-stomatal-conductance-and-transpiration/minimum-epidermal-conductance-gmin-a-k-a-cuticular-conductance/. For samples that displayed non-linear change in mass over time, we discarded the initial 1-2 measurements under the assumption that these were most likely to be artificially high due to incomplete stomatal closure. *g*_*min*_ was calculated from the remaining measurements by fitting a linear regression to the relationship between mass change and the atmospheric vapor pressure deficit. In most cases, one flower and leaf per plant was sampled, and previous measurements suggest that intraspecific variation in flower *g*_*min*_ is relatively small (Roddy *et al*., 2016). Leaf (*LMA*) or petal mass per area (*PMA*) were quantified from the area and dry mass measurements made on samples measured for *g*_*min*_. Water content per area (*W*_*area*_) was calculated as the difference between the initial mass and dry mass divided by the surface area of each sample, and the water content per dry mass (*W*_*mass*_) was calculated as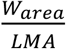 converted to units of grams of water per gram of dry mass. Water residence time (τ) was calculated as 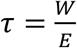 where *E* is the transpiration rate, which was calculated from *g*_*min*_ assuming a vapor pressure deficit of 1 kPa (Simonin *et al*., 2013; Roddy *et al*., 2018). Using a higher vapor pressure deficit reduces τ.

Most of the vein density (*D*_*v*_) data have been published previously (Roddy *et al*., 2013) and sampling methods are briefly summarized here. For leaves, approximately 1-cm^2^ sections from midway between the leaf midrib and margin, midway between the base and tip of the leaf were excised and immediately placed into 4% NaOH. To account for the high variability in vein density within a petal, we collected multiple 1-cm^2^ sections from throughout the petals and placed them in 4% NaOH. After 2-4 weeks, leaves were washed in distilled H_2_O, transferred to a 3% bleach solution for approximately 20 minutes, washed again in distilled H_2_O, and then placed in 95% ethanol. After clearing in NaOH, petals were washed in distilled H_2_O and transferred into ethanol, skipping the bleaching step. Once in ethanol, samples were briefly stained with Safranin O and imaged at 5-20x magnification under a compound microscope outfitted with a digital camera. One or two images per section from each of five to twelve sections per species were captured. For each image, the total length of veins was measured manually using ImageJ [version 1.44o; Schneider *et al*. (2012)] and divided by the total area of the image to calculate vein length per area. The mean for each structure of each species was calculated and used for subsequent analyses.

### Data analysis

All analyses were conducted in R (v. 4.1.2) using log-transformed data. We removed species with outlying data for any trait, which were a total of six species across all traits. A broadly inclusive, dated phylogeny was created using V.Phylomaker (Jin & Qian, 2019) after looking up higher level family classification using the R package *taxize* (Chamberlain & Szocs, 2013).

Differences between organs in each trait were determined using paired t-tests. Phylogenetically controlled paired t-tests were conducted using the function *phyl*.*pairedttest* in the R package *phytools* (Revell, 2012). Standard major axis regressions and their slope and elevation tests were conducted using the R package *smatr* (Warton *et al*., 2012).

## Notes

### Competing Interest Statement

The authors have declared no competing interest.

